# Multi-omic surveillance of *Escherichia coli* and *Klebsiella* spp. in hospital sink drains and patients

**DOI:** 10.1101/2020.02.19.952366

**Authors:** B Constantinides, KK Chau, TP Quan, G Rodger, M Andersson, K Jeffery, S Lipworth, HS Gweon, A Peniket, G Pike, J Millo, M Byukusenge, M Holdaway, C Gibbons, AJ Mathers, DW Crook, TEA Peto, AS Walker, N Stoesser

## Abstract

*Escherichia coli* and *Klebsiella* spp. are important human pathogens that cause a wide spectrum of clinical disease. In healthcare settings, sinks and other wastewater sites have been shown to be reservoirs of antimicrobial-resistant *E. coli* and *Klebsiella* spp., particularly in the context of outbreaks of resistant strains amongst patients. Without focusing exclusively on resistance markers or a clinical outbreak, we demonstrate that many hospital sink drains are abundantly and persistently colonised with diverse populations of *E. coli, Klebsiella pneumoniae* and *Klebsiella oxytoca*, including both antimicrobial-resistant and susceptible strains. Using whole genome sequencing (WGS) of 439 isolates, we show that environmental bacterial populations are largely structured by ward and sink, with only a handful of lineages, such as *E. coli* ST635, being widely distributed, suggesting different prevailing ecologies which may vary as a result of different inputs and selection pressures. WGS of 46 contemporaneous patient isolates identified one (2%; 95% CI 0.05-11%) *E. coli* urine infection-associated isolate with high similarity to a prior sink isolate, suggesting that sinks may contribute to up to 10% of infections caused by these organisms in patients on the ward over the same timeframe. Using metagenomics from 20 sink-timepoints, we show that sinks also harbour many clinically relevant antimicrobial resistance genes including bla_CTX-M_, bla_SHV_ and *mcr*, and may act as niches for the exchange and amplification of these genes. Our study reinforces the potential role of sinks in contributing to Enterobacterales infection and antimicrobial resistance in hospital patients, something that could be amenable to intervention.

**IMPORTANCE:** *Escherichia coli* and *Klebsiella* spp. cause a wide range of bacterial infections, including bloodstream, urine and lung infections. Previous studies have shown that sink drains in hospitals may be part of transmission chains in outbreaks of antimicrobial-resistant *E. coli* and *Klebsiella* spp., leading to colonisation and clinical disease in patients. We show that even in non-outbreak settings, contamination of sink drains by these bacteria is common across hospital wards, and that many antimicrobial resistance genes can be found and potentially exchanged in these sink drain sites. Our findings demonstrate that the colonisation of handwashing sink drains by these bacteria in hospitals is likely contributing to some infections in patients, and that additional work is needed to further quantify this risk, and to consider appropriate mitigating interventions.

## INTRODUCTION

Infections caused by Enterobacterales, including *Escherichia coli* and *Klebsiella* spp., are major causes of global morbidity, and particular antimicrobial-resistant strains (namely extended-spectrum beta-lactamase and carbapenemase producers) have been listed as critical priority pathogens for mitigation by the WHO. In the UK, year-on-year increases have been observed in the number of *E. coli* and *Klebsiella* spp. bloodstream infections (1), for reasons which remain unclear. As well as causing invasive disease, these organisms are capable of colonising a wide range of animal and environmental niches, and are frequently carried in the human gastrointestinal tract (2). As such, they are also commonly found in human wastewater, and in wastewater-associated sites such as sewers and water treatment infrastructure (3).

A significant proportion of Enterobacterales infections are healthcare-associated, prompting the UK government to introduce a target in 2016 to halve the number of healthcare-associated Gram-negative bloodstream infections by 2021 (4). Wastewater sites in hospitals have been highlighted as reservoirs of drug-resistant Enterobacterales, with several studies reporting that ongoing transmission and outbreaks of human disease are associated with the contamination of, for example, sinks, by these organisms (5, 6). More recently, several studies have shown reductions in colonisation and/or invasive infection with Enterobacterales and other Gram-negative bacilli following the introduction of strategies to remove sinks and mitigate possible contamination from wastewater sources in patient rooms (7, 8). Most of these studies however focus on the sampling and control of antimicrobial-resistant strains, often representing a more immediate clinical problem in an outbreak setting, rather than on the possibility that these sites may represent part of the wider endemic transmission network of both susceptible and resistant strains causing infection in patients.

Whole genome sequencing of bacterial isolates is increasingly used as the most robust, high-resolution approach to characterising relatedness between strains, and hence determining likely transmission (9). However, the diversity of complex, polymicrobial environmental reservoirs is incompletely captured by sequencing small numbers of isolates, and this breadth of diversity can be more fully captured by using a metagenomic approach, which characterises the genetic complement of a whole sample (10). Combining both approaches has been shown to improve our understanding of species and antimicrobial resistance (AMR) gene diversity within environmental, wastewater and river samples (11, 12) and of transmission in a sink-associated outbreak of *Sphingomonas koreensis* (also a Gram-negative bacillus) in the NIH Clinical Centre in the US (13).

In order to investigate the prevalence of contamination of healthcare sinks by strains of *E. coli* and *Klebsiella* spp., including those resistant to third-generation cephalosporins and carbapenems, we sampled all sink sites using p-trap (U-bend) aspirates across several wards and timepoints in a single UK hospital in 2017. We used a combination of whole genome sequencing of cultured isolates from all sink samples and metagenomic sequencing of a subset of sink samples to facilitate a high-resolution assessment of the genetic diversity present in these niches. To determine whether sinks were a reservoir of Enterobacterales strains causing infection in patients over similar timeframes, we simultaneously retrieved relevant isolates from culture-positive specimens taken from patients admitted to the same ward locations, and used genomics to identify the degree of genetic relatedness.

## RESULTS

### Diverse, often antimicrobial-resistant Enterobacterales strains are frequent—and often persistent—colonisers of hospital sink drains

439 Enterobacterales isolates comprising *E. coli* (n=180), *K. oxytoca* (n=166) and *K. pneumoniae* (n=93) were cultured and successfully sequenced at one or more timepoints from 12/20 (60%), 9/23 (43%) and 16/23 (70%) sinks sampled four times over 12 weeks (March-May 2017) in general medicine (GM), adult critical care (ACC) and acute admissions (AA) wards respectively (97/264 [37%] sink-timepoints culture-positive overall; Figure 1). A further 30 isolates of *E. coli* (n=13), *K. oxytoca* (n=13) and *K. pneumoniae* (n=4) were cultured from 11/59 (19%) sinks in a haematology ward, sampled at a single timepoint only during this period (Figure S1). See Table S1 for surveyed sink descriptions. Species distributions (by culture) were relatively even across the general medicine ward, while the adult critical care unit was enriched for *E. coli*, and the acute admissions ward was depleted in *K. pneumoniae* (Table S2).

**FIG 1.**
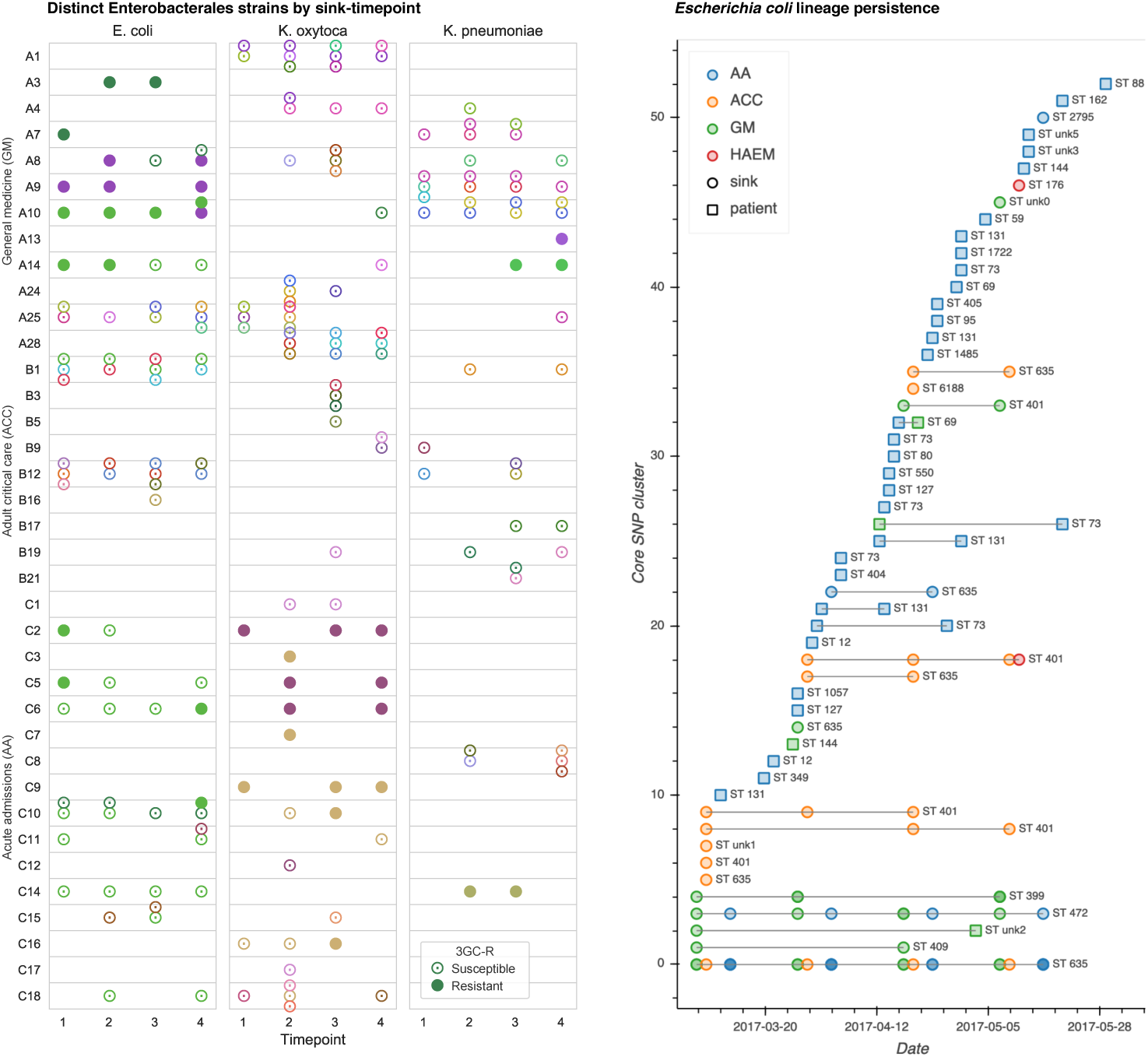
Cluster distribution and persistence. Left: strain-distinct cultured isolates of *E. coli, K. oxytoca* and *K. pneumoniae* from sink drain aspirates sampled over twelve weeks across three hospital wards. Different colours indicate distinct strains (defined by 100 SNP clusters), and cefpodoxime-resistant and/or selected ESBL-positive isolates are indicated by filled markers. Right: persistence of sink and contemporaneous patient *E. coli* strains throughout the sampling period.

Analysis of whole genome sequences from cultured Enterobacterales revealed widespread and sustained colonisation of sinks by multiple sequence types (STs) of these species (Figure 1, Figure S1) In total, 8 known and 4 novel *E. coli* STs were represented, of which STs 635 (n=109, 61%), 401 (n=25, 14%) and 472 (n=18, 10%) accounted for 84% of sequenced isolates (152/180). *Klebsiella* spp. STs were more varied: 15 known and6 novel *K. oxytoca* STs were represented, of which the most frequent was ST177 (33/166, 20%), while there were 18 known and 1 novel *K. pneumoniae* STs, the most frequent being ST872 (24/93, 26%).

Across all locations and sink-timepoints, sequenced isolates comprised 20, 50 and 26 distinct strains (defined as differing by ≤100 recombination-adjusted core SNPs; see Materials and Methods) of *E. coli, K. oxytoca* and *K. pneumoniae* respectively (Figure 1, Table 1). Positive sinks cultured up to three of these distinct strains per species at any timepoint (Figure 1, Figure S1), reflecting significant diversity within species in sink niches. Of the 37 longitudinally sampled culture-positive sinks from which sequences were obtained, 31 (84%) grew isolates belonging to the same strain across multiple timepoints, highlighting persistent background colonisation illustrated for *E. coli* and *Klebsiella* spp. in respective figures 1 and S5. Isolates resistant to third-generation cephalosporins were cultured at 16 sink-timepoints across 12 distinct sinks, with resistant and susceptible cultures of the same genetic strain co-occurring in 11/16 (69%) sink-timepoints, suggestive of gain and/or loss of genes conferring cephalosporin resistance in this setting. No carbapenem-resistant isolates were cultured.

**TABLE 1.** Spatiotemporal distribution of 100 core SNP lineages of cultured *E. coli* (n=53), *K. oxytoca* (n=51) and *K. pneumoniae* n=30) *A)* overall and *B)* occurring in >1 isolate in sinks on wards that were repeatedly sampled.

| A) |  | *E.coli* | *K. oxytoca* | *K. pneumoniae* | Total |
| --- | --- | --- | --- | --- | --- |
| **Lineages with 1 isolate** |  |  |  |  |  |
|  | Patient | 28 | 1 | 3 | 32 |
|  | Sink | 3 | 37 | 12 | 52 |
| **Lineages with >1 isolate** |  |  |  |  |  |
| Single timepoint | Same sink | 5 | 6 | 3 | 14 |
|  | Different sinks; same ward | 0 | 1 | 0 | 1 |
| Multiple timepoints | Same sink | 7 | 1 | 7 | 15 |
|  | Different sinks; same ward | 1 | 4 | 4 | 9 |
|  | Different wards | 3 | 1 | 0 | 4 |
| Patients | Patient and sink (single timepoint) | 1* | 0 | 0 | 1 |
|  | Same patient | 1 | 0 | 1 | 2 |
|  | Different patients | 4 | 0 | 0 | 4 |
| Total |  | 53 | 51 | 30 | 134 |

\*lineage has 3 isolates; all taken from the same ward; 2 from the same sink at the same timepoint and 1 from a patient 2 months later.

| B) |  | *E.coli* | *K. oxytoca* | *K. pneumoniae* | Total |
| --- | --- | --- | --- | --- | --- |
| **Sink lineages with >1 isolate (excludes haematology ward)** |  |  |  |  |  |
| Single timepoint | Same sink | 5 | 4 | 3 | 12 |
|  | Different sinks; same ward | 0 | 0 | 0 | 0 |
| Multiple timepoints | Same sink | 8 | 1 | 7 | 16 |
|  | Different sinks; same ward | 1 | 4 | 4 | 9 |
|  | Different wards | 2 | 1 | 0 | 3 |
| Total |  | 16 | 10 | 14 | 40 |

### Enterobacterales can be highly abundant in sink drains, representing dominant populations in some wards

Deep metagenomic Illumina sequencing was performed for 20 sink-timepoints on p-trap aspirates from seven sinks on the three wards at two timepoints, and all four timepoints for a single sink unit in the adult critical care ward (median 3.6m reads/sample; IQR: 3.3m-7.2m). The three most abundant bacterial genera were *Klebsiella, Escherichia* and *Citrobacter*, all common healthcare-associated pathogens (Figure 2). Sink drains in the general medicine ward were the most abundantly colonised by Enterobacterales (Figure 2), to which more than 50% of reads were assigned, and were markedly less diverse than those in adult critical care and acute admissions wards, which had a dominance of *Klebsiella* spp., mirroring the culture results. 90% of species-level classifications in the sink-timepoints from the general medicine ward came from a median of just 21 bacterial species, compared with medians of 310 and 450 species in the adult critical care and acute admissions wards respectively. Microbial composition varied markedly between sampling timepoints for individual sinks, but sinks within wards exhibited more similar taxonomic profiles than those between wards (Figure 2), suggesting distinct ward-based wastewater ecologies. Total metagenomic sequence content was hierarchically structured by ward and by sink (Figure S2). Staff room sink A25 exhibited distinctive taxonomic and *k*-mer profiles from patient room sinks in the general medicine ward (Figure 2, Figure S2).

**FIG 2.**
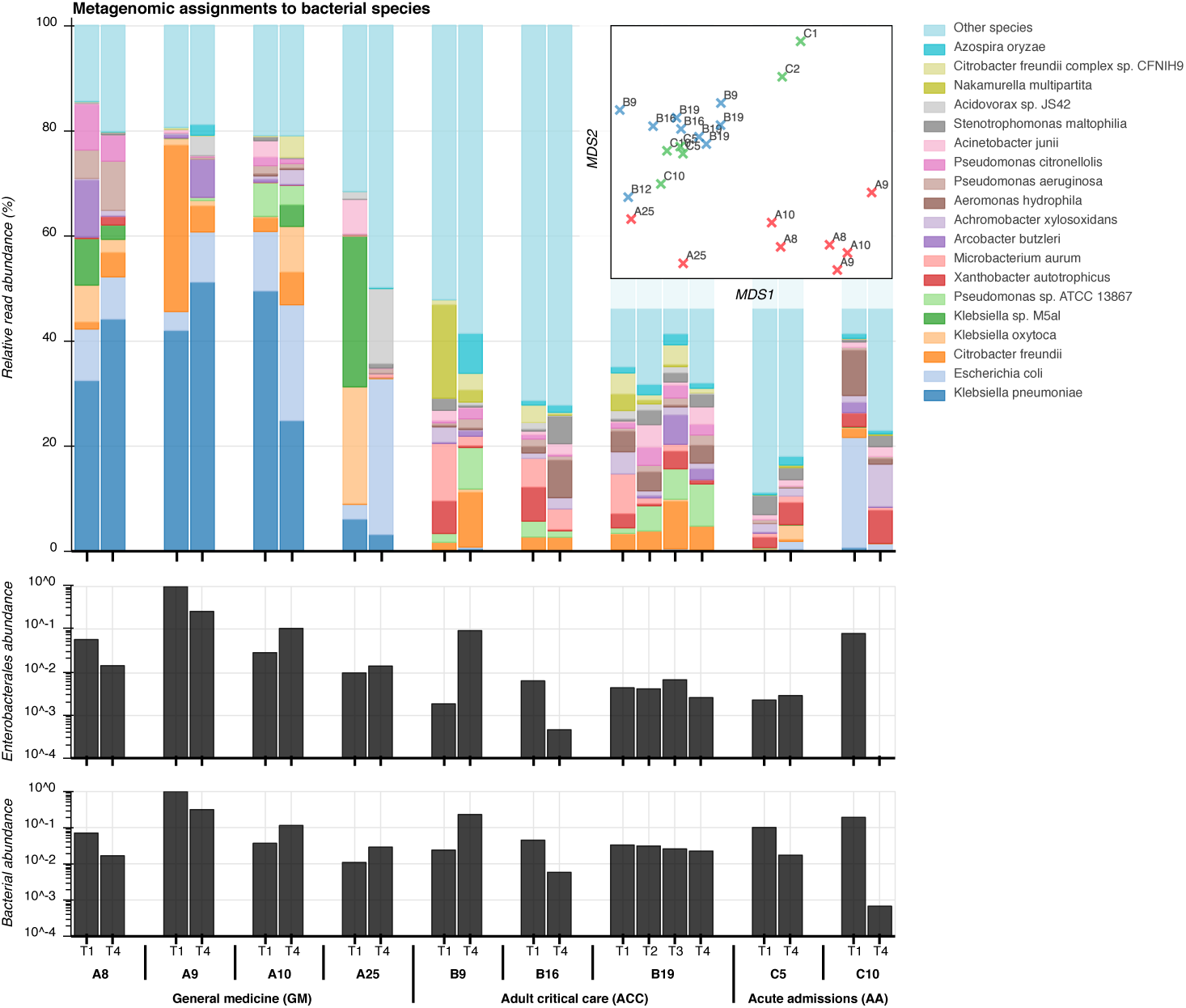
Taxonomic composition of sink microbiota from metagenomic sequencing. Top: relative abundance of the 20 most abundant bacterial species among sink drain aspirates (Kraken), inset with a corresponding multidimensional scaling (MDS) projection of pairwise distances between samples. Centre: spike-normalised relative abundance of species classifications at or below the order Enterobacterales among sink-timepoints. Bottom: spike-normalised relative abundance of Kraken classifications at or below the superkingdom Bacteria.

Sinks with high metagenomic abundance of the three Enterobacterales species reliably yielded corresponding cultures. The area beneath the receiver operating characteristic (ROC) curve for culture-based detection of these species was 0.93 (Figure S3). When the relative metagenomic abundance of a species was above 0.1%, 1% and 10%, one or more cultures of the same organism were obtained in 58% (18/31), 76% (16/21) and 89% (8/9) of sinks respectively. Conversely, culture detection therefore failed in 42% (13/31), 24% (5/21) and 11% (1/9) of cases where an Enterobacterales species was present at or above respective thresholds of 0.1%, 1% and 10% metagenomic abundance. A single sink-timepoint (first sample from A8; general medicine) failed to culture any Enterobacterales, but yielded 4%, 5% and 16% relative metagenomic abundances for the three study species. Thus metagenomic sequencing suggests that persistence may be even more widespread than indicated by culture alone.

### Most environmental Enterobacterales appear to cluster within specific sinks and wards, except for *E. coli* ST635, which is widely distributed

The 96 strains found in sinks exhibited structure at the ward and sink level. Four strains were found in multiple wards (predominantly *E. coli* ST 635, but also STs 472, 401 and *K. oxytoca* ST 146). 92 (96%) strains were only ever found in a single ward, and of the 44 strains cultured twice or more, only 14 (32%) were cultured from different sinks (Table 1A). Further, of the 40 strains cultured twice or more on wards which were repeatedly sampled (i.e. excluding the haematology ward which was only sampled once), 12 (30%) were only seen at the same sink-timepoint, 16 (40%) were seen in the same sink at different timepoints, and 12 (30%) in different sinks at different timepoints (1B). This structure was reflected in the recombination-adjusted core genome species phylogenies (Figure 3). The main exception to ward and sink-based clustering was *E. coli* ST635, which comprised more than half of isolates sequenced from sinks, and was found in 13/20 (65%) *E. coli*-positive sinks.

**FIG 3.**
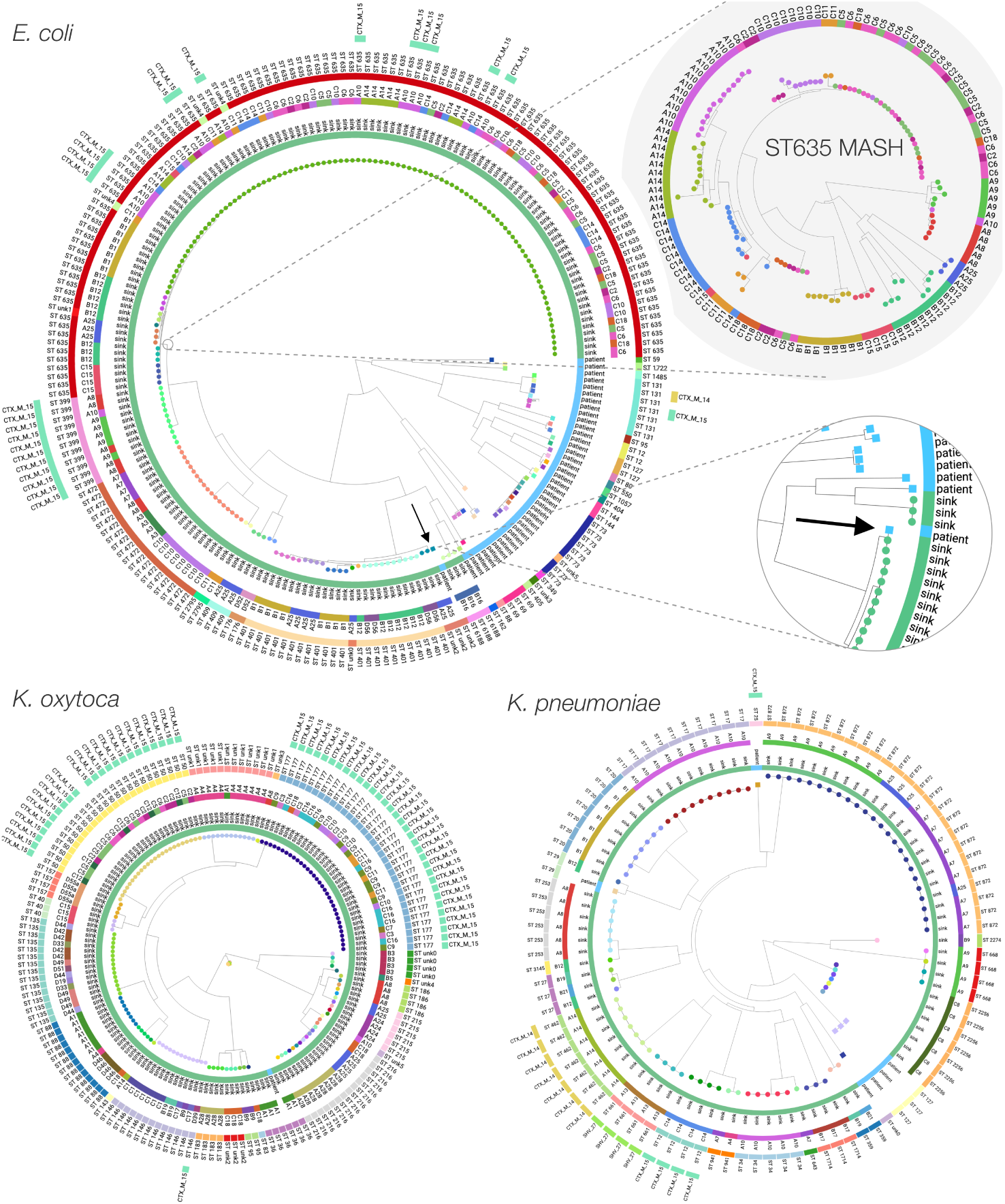
Maximum likelihood phylogenies of *E. coli, K. oxytoca* and *K. pneumoniae* cultured from sink drain aspirates sampled over twelve weeks across three wards, with two zooms corresponding to an *E. coli* ST635 neighbour-joining MASH subtree whose tips are coloured by sink, and genetic overlap between a sink culture and a urine culture from a patient with ward contact during the study. Tip colours indicate strains, with rings inside-to-out denoting: patient/sink, sink designation, sequence type, and ESBL genotype.

However, there was sink-level clustering even within *E. coli* ST635, more clearly shown in neighbour-joining trees constructed from pairwise read-based MASH distances, representing both core and accessory genomic content (see *E. coli* ST635 zoom in Figure 3; colours indicate distinct sinks). Although pairwise correlations between core and accessory genomic distances were high (Table S3), incorporating accessory content yielded additional resolution beyond core SNP distances (Figure S4). Permutational analysis of variance (PERMANOVA) using pairwise core SNP and read-based MASH distances supported significant grouping of isolates from all three species by ward and by sink (P<0.001), most conclusively for *K. pneumoniae* (Table S4).

### Patient *E. coli* isolates were more diverse than those found in sinks, and included isolates from known ‘high-risk’ clinical lineages

From March to May 2017, 1384 relevant clinical samples from 719 patients were submitted to the microbiology laboratory for processing (AA n=779, ACC n=365, GM n=240), of which 397/1384 (29%) were positive for microorganisms, and 107 were culture-positive for one of the study organisms (*E. coli* [n=96], *K. oxytoca* [n=2], *K. pneumoniae* [n=9]). 46/107 (43%) of these isolates were retrieved for sequencing, including 19/22 isolates from bloodstream infections, 3/6 from respiratory samples, 21/73 from urine samples, and 0/10 other samples.

Among 39 sequenced *E. coli* patient isolates, 21 STs were represented, including known high-risk lineages (23/39 (59%) isolates) not seen in sinks: namely ST73 (n=8), ST131 (n=7), ST69 (n=3), ST12 (n=2), ST127 (n=2), and ST95 (n=1). The single sequenced *K. oxytoca* isolate was ST36, and the six *K. pneumoniae* isolates came from four STs, including two high-risk lineages, ST25 and ST29. Across the three species, there were 34, 1 and 4 distinct lineages, respectively (Table 1).

### Genetic similarity of sequenced patient and environmental isolates

As well as being diverse, the 39 clinical *E. coli* isolates were phylogenetically distinct from most sink isolates, which largely came from just four sequence types (ST635, ST401, ST472 and ST399). The exception was an *E. coli* isolated from urine taken on the general medicine ward, which was 17 and 19 core SNPs from two isolates from sink A25 in the same ward, sampled 58 days prior to the clinical sample (Figure 1; right; cluster 4). A read-based MASH distance of 7 × 10^−6^ between this pair of isolates indicated very high total genomic (chromosome+accessory) similarity. A records search for admissions of this patient prior to commencing sink sampling revealed four inpatient admissions (of 0 (day case), 1, 2 and 5 nights’ duration), of which 3 included time on the acute admissions ward. There were no prior admissions onto the general medicine ward (in which their positive urine specimen was taken), although their eleven-night spell on the general medicine ward commenced with a seven-hour episode in acute admissions. The positive clinical specimen was taken ten days after the patient’s admission onto the general medicine ward, indicating a large duration of exposure to a ward environment shown to be harbouring a very similar strain of *E. coli* to the patient’s urine culture. The next most closely related *E. coli* clinical isolate was 3,688 core SNPs from its nearest sink neighbour (read-based MASH distance 0.009), reflecting the otherwise large evolutionary distances separating the cultured clinical and environmental *E. coli* (Figure 3).

Unlike *E. coli*, the small numbers of *Klebsiella* spp. patient isolates were not phylogenetically segregated from environmental isolates, but the closest patient and sink isolates differed by 2,558 core SNPs, indicating a lack of observed overlap over these timeframes.

### Antimicrobial resistance genes are prevalent and spatially structured in sinks

The presence of 571 clustered CARD antimicrobial resistance genes in cultured isolates was supported by ≥75% exactly matching read coverage reported by ResPipe (Figure 4). Among these were known transmissible genes of clinical concern including beta lactamases (e.g. bla_TEM_, bla_CTX-M_, bla_SHV_), aminoglycoside resistance genes (*aac(3), aac(6)* families) and quinolone resistance genes (*qnr* family). Some of these, including cmlA and qacH, were widely seen in sink metagenomes but less frequently in cultured isolates, consistent with a background resistance reservoir that may pose a risk in different populations to those cultured (of either same or different species). Spatial structure was evident among both cultured isolates and metagenomes, although resistance repertoires of isolates frequently clustered across ward boundaries, in keeping with findings of our prior core genome analysis. Resistance genes detected in cultured sink isolates were also abundant within sink metagenomes at one or more timepoints. Sink drain metagenomes yielded 673 CARD genes exceeding 75% coverage, and after clustering large gene families represented by many similar sequences (see methods for detailed description), only five genes abundant in one or more cultured isolates were not detected in at least one metagenome. Notably, these five genes (*gadW, len-26, tet(B), mgrA* and *sat-2*) were all seen in isolates from sinks not subject to metagenomic sequencing, showing that resistance genes cultured from sink drains were highly contained in corresponding metagenomes.

**FIG 4.**
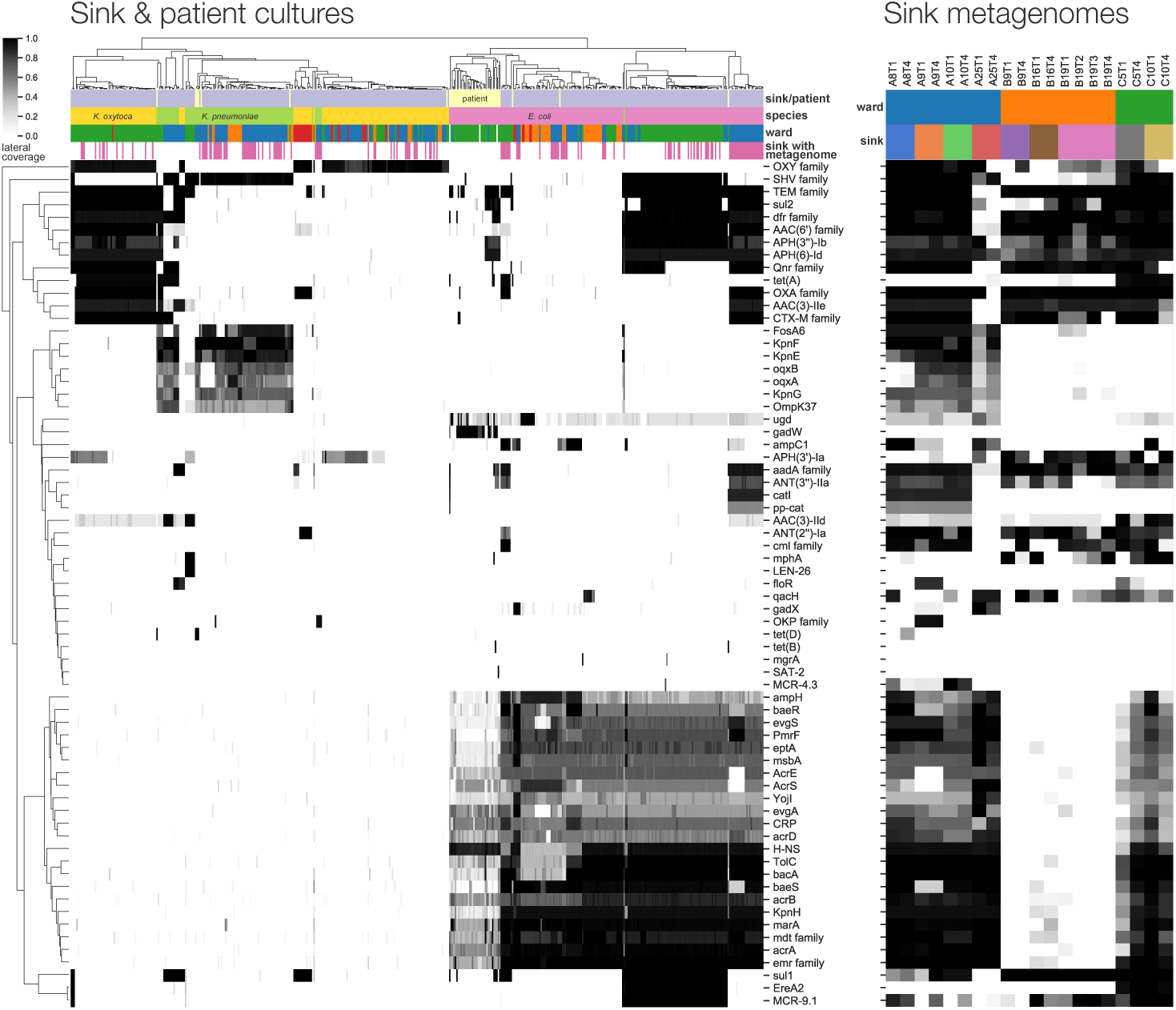
Antimicrobial resistance gene content of cultured isolates and sink drain metagenomes. Left: lateral coverage of ResPipe/CARD genes within sink drain and clinical isolates. Displayed genes attained ≥75% lateral coverage in one or more isolates. Right: Corresponding lateral coverage of the same genes in sink drain aspirate metagenomes.

Third generation cephalosporin-resistant phenotypes in Enterobacterales sink isolates could be explained by the presence of major extended-spectrum beta-lactamase (ESBL) genes blaSHV-27, bla_CTX-M-14_ and bla_CTX-M-15_, detected by ARIBA/CARD in 4, 8 and 87 isolates respectively. bla_CTX-M-15_ was identified in two distinct strains of *E. coli* ST635 and ST399, restricted to three bay sinks (A8, A9, A10) in three adjacent rooms of the general medicine ward. bla_CTX-M-15_-positive *K. oxytoca* ST50 and ST177 were identified in 10 sinks (C2-3, C5-6, C7, C9-12, C16) on the acute admissions ward. bla_SHV-27_, bla_CTX-M-14_ and bla_CTX-M-15_ were observed in *K. pneumoniae* from sinks A13, A14 (general medicine) and C14 (acute admissions) respectively; one *K. pneumoniae* patient isolate was also bla_CTX-M-15_-positive. These findings suggest sink-associated isolates, such as *E. coli*, may represent reservoirs of clinically relevant resistance genes.

Surprisingly, the colistin resistance gene *mcr-4* was detected in the metagenomes—yet not cultured genomes—of three sinks in adjacent bays of the general medicine ward (A8-10). Assembly of the sink A10 metagenome generated a 5.4kbp plasmid sequence containing an *mcr-4* gene with 98.8% overall identity at 94% query coverage to an 8.7kbp pMCR-4.2 plasmid previously reported in pigs from Italy, Spain and Belgium (14). This *mcr* variant has been previously reported in European Acinetobacter, Enterobacter, Salmonella, and Escherichia spp. but not to our knowledge in the United Kingdom. Screening all metagenomes for assembled *mcr-4* produced alignments in two sinks on the ward (A8, A9) across a total of six sink-timepoints, with coverage and abundance suggesting low and declining prevalence of this gene over time (Table S5). An *mcr*-positive *E. coli* (reported as *mcr-4*.*3*) was cultured from sink C5 (acute admissions; second timepoint) according to both ARIBA/CARD and ResPipe/CARD. Metagenomic sequencing was performed for the first and the fourth but not the second timepoint aspirate for this particular sink. Another *mcr* gene, *mcr-9* was more widespread, and detected with complete coverage in 77 cultured isolates across 11 distinct sinks, predominantly but not exclusively in *E. coli* (73/77 occurrences).

### Metagenomic screening suggests that clinical isolates may be more widely present in the environmental reservoir than observed from culture-based comparisons

Sink metagenomes were individually screened for *k*-mer containment of *i)* strain-representative sink isolate genome assemblies, *ii)* strain-representative patient isolate genome assemblies and *iii)* core genomes of selected control organisms, including five clinical core genomes each from pathogenic strains of *E. coli* and *Klebsiella* spp. from Bush et al. (15), together with NCBI canonical species references for several pathogens expected to be absent from sink drain microbiota (Figure 5; left). This demonstrated similar sink, ward, and temporal structure to that of culture, particularly underlying similarities in the microbiota of nearby sinks, as well as flux between sampling timepoints. Strain-representative sink culture assemblies from the same sink and timepoint as the screened sink metagenome were the best contained, sharing the most *k*-mer hashes. Assembled isolates originating from the same sink but at a different timepoint to the screened metagenome shared significantly fewer *k*-mer hashes (P=0.013) than same sink/same timepoint comparisons. The containment of strain-representative assemblies from different sinks in the same ward as the metagenome was significantly lower still (P<0.0001), and so in turn were the remaining comparisons of cultures grown from different sinks in different wards to the screened metagenome (P<0.0001).

**FIG 5.**
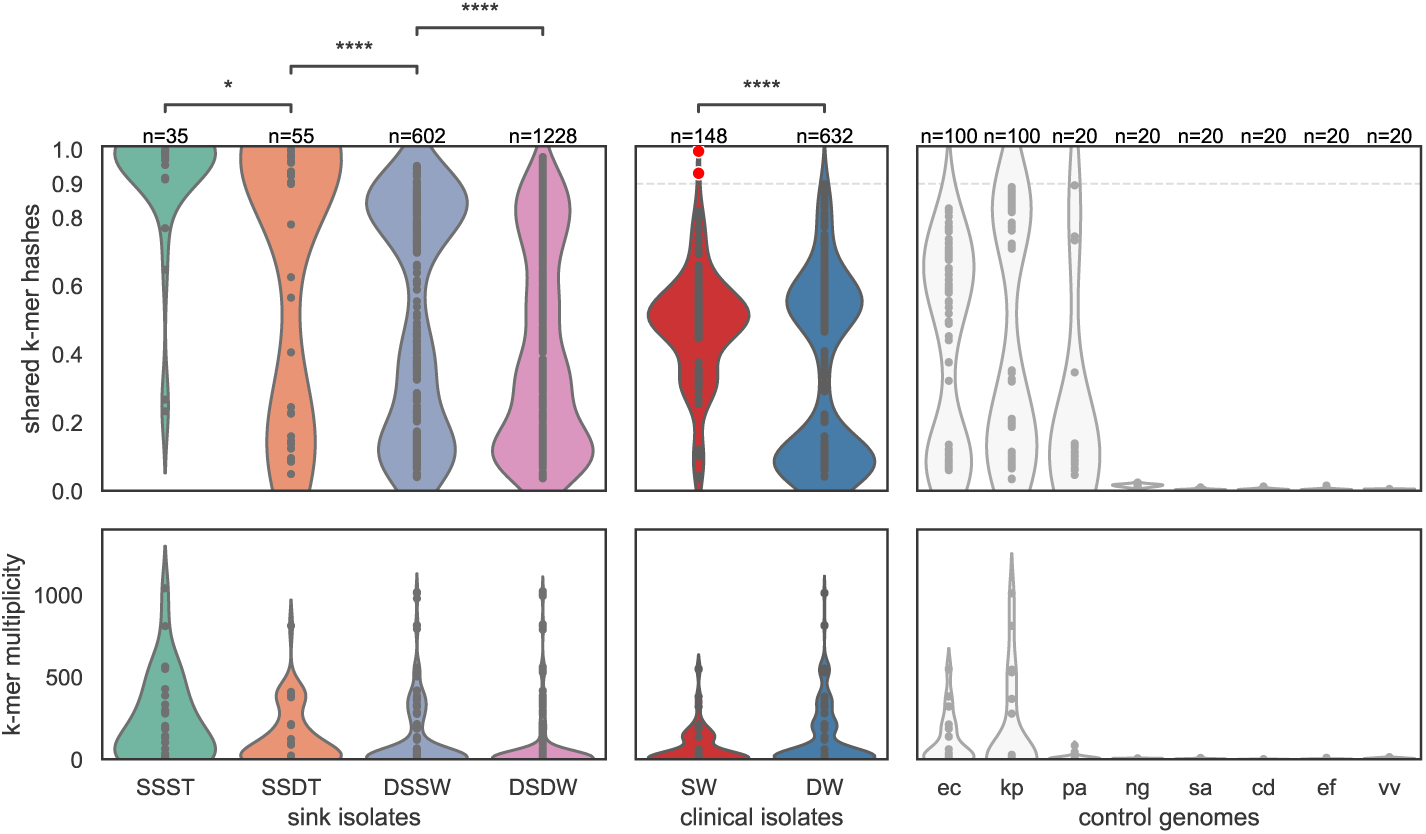
Metagenomic containment of sink (left) and patient (centre) cultured strain-representative genome assemblies, and control genomes (right). Shared *k*-mer hashes and median *k*-mer multiplicity values are as reported by MASH Screen. SSST=same sink and same timepoint; SSDT=same sink at different timepoints (shared hashes Mann-Whitney U P=0.013 vs. SSST); DSSW=different sinks of the same ward (P<0.0001 vs. SSDT); DSDW=sinks on a different ward (P<0.0001 vs. DSSW). SW=strain-representative assemblies of clinical isolates in the same ward; DW=strain-representative assemblies of clinical isolates from a different ward (P<0.0001 vs. SW). Control genomes comprised *E. coli, K. pneumoniae P. aeruginosa, N. gonorrhoeae, S. aureus, C. difficile, E. faecalis*, and *V. vulnificus*, shown abbreviated with binomial initials. A case of within-ward sink-patient overlap is highlighted with red markers, corresponding to high strain containment in the metagenomes of sink A25 timepoints 1 and 4.

Among control genomes, *k*-mer hashes shared between sink metagenomes and the core genomes of *Neisseria gonorrhoeae, Staphylococcus aureus, Clostridioides difficile, Enterococcus faecalis* and *Vibrio vulnificus*did not exceed 3%. Reference genomes of *E. coli, K. pneumoniae* and *Pseudomonas aeruginosa* were abundant and highly contained by many metagenomes, but none exceeded 90% shared *k*-mer hashes (Figure 5; right).

In contrast, screening for strain-representative patient assemblies in sink metagenomes revealed significantly greater similarity (P<0.0001) between patient and environmental Enterobacterales strains collected from the same ward than from different wards (Figure 5; centre), supporting genetic overlap between clinical isolates from patients and uncultured isolates in sink niches in a given ward setting. Indeed, the only strain-representative patient isolate with greater than 90% sink metagenome *k*-mer containment was the *E. coli* urine culture described in the aforementioned case of sink-patient overlap, of which 99.5% and 93.0% of *k*-mers were contained within the respective A25T1 and A25T4 sink metagenomes (Figure 5; centre; red markers).

## DISCUSSION

In this study, we have demonstrated that hospital sink drains are widely—and in many cases abundantly—contaminated with key Enterobacterales species causing healthcare-associated infections, and are potential reservoirs of multiple resistance genes encoding resistance to important clinical antimicrobials. Populations of antimicrobial-resistant and susceptible *E. coli* and *Klebsiella* spp. may be persistent colonisers of sinks, and different wards may have markedly different sink ecosystems, perhaps reflecting different and potentially modifiable infrastructures, selection pressures, and contributing sources. Ward and sink-level genetic structure was most evident within the accessory genome, and observed repertoires of transmissible resistance genes often transcended species boundaries, instead clustering more tightly by sink unit. Characterising these highly diverse reservoirs is difficult, and we have shown that combination approaches utilising metagenomics and sequencing of cultured isolates are complementary in understanding the diversity of species, strains, and the resistance genes present within these niches. For example, metagenomics highlighted several cases of abundant *mcr-4* in sink drain aspirates from which cultured Enterobacterales isolates did not carry the gene.

Colonisation patterns of sink niches differed markedly between the two genera investigated. *E. coli* strains have evolved to colonize and adapt to multiple niches, including some which have adopted pathogenic lifestyles, and appear to have different distributions in humans, domesticated and wild animals, and the environment. There is however no absolute correlation between phylogenetic lineage and any given niche, and overlaps are observed. Interestingly, in our study, more than half of the *E. coli* sink isolates cultured were ST635, which has been recently described as a highly adapted, resistance- and virulence gene-enriched wastewater-associated strain thought to be globally distributed, but is also found in humans, animals and other environments (16). Of note, it has been observed in association with several clinically relevant transmissible resistance genes, including ESBLs, carbapenemases, and rRNA methylases, and was one of only two *E. coli* STs in our study that harboured an ESBL (bla_CTX-M-15_). We observed presence/absence of bla_CTX-M-15_ across closely related ST635 isolates, suggesting that this gene may be frequently lost/gained in sinks. Also notable in the context of ST635 was the ability of read-based *k*-mer composition to resolve fine-grained structure between the populations of different sinks, beyond that observed in the core-only SNP phylogeny. Other common *E. coli* sink lineages were ST399 and ST472, which to date have predominantly been seen in humans/animals, rather than the environment.

The phylogenetic distribution of sink isolates of *K. pneumoniae* appeared to mirror that seen in a global collection of isolates, providing little evidence that a particular lineage was predominating in, or particularly adapted to, the wastewater environment. Studies of the population structure of unselected *K. oxytoca* are limited, but again we observed a diverse population amongst sink isolates, with a deep branch separating two distinct groups as previously described. Interestingly, two *K. oxytoca* strains associated with bla_CTX-M-15_ were widely distributed amongst sinks in the acute admissions ward; outbreaks of ESBL- and carbapenem-associated *K. oxytoca* in association with contaminated handwashing sinks have been described in other settings (17).

Genomic overlap with sink isolates was identified in 1/46 (2%; 95% CI: 0.05-11%) of all sequenced isolates causing clinical infections over the same timeframe, with a temporal association consistent with acquisition from a sink source (i.e. sink isolate observed first), and following ten days of patient exposure to a ward environment wherein the overlapping strain was previously cultured. We may have significantly underestimated the degree of overlap between these two compartments for several reasons. Firstly, we have shown the diversity in sink niches is substantial, and with a culture-based approach agnostic to any selective marker, even sequencing 444 isolates from 48 sinks will have limited ability to capture the underlying diversity for complete comparison of sink-patient pairs at the isolate-level. Supporting this, screening the metagenomes of a subset of 20 sinks using patient isolates suggests that overlap between these reservoirs may be more common than observed at the isolate-level. Second, clinical isolates represent the tip of the iceberg of any transmission chain, with the majority of transmission events likely occurring between gastrointestinal tract (asymptomatic carriage) and the wastewater environment. Nonetheless, in the context of understanding how sinks may be contributing to infection caused by *Klebsiella* spp. and *E. coli*, focusing on clinical isolates seems appropriate. Third, the interval between sampling dates for our observed patient-sink isolate-pair was 58 days, suggesting that the timeframe between acquisition from the environment and infection may be long, and may not be adequately captured with a study timeframe spanning three months. In addition, a major study limitation is the fact that only 46/107 patient isolates could be successfully retrieved (due to the high turnover of samples in our high-volume service laboratory), and *Klebsiella* spp. cultures were especially limited. The risks of transmission and possibly sink-associated infection could be more clearly defined by more extensive sampling over a greater timeframe, and thorough investigation into the exchange of resistance-associated mobile genetic elements, but would require a considerable increment in resource. Characterising microbial diversity present on sink strainers would also be of benefit, as the risks of droplet-mediated dispersal from sink drains have been shown to be most significant when the sink drain is located immediately below the tap, and if the organisms migrate from the sink trap onto the strainer (18, 19). However, given the different sink structures across wards, the p-trap was the only site which could be consistently sampled (since ACC had horizontally draining sinks without strainers). Characterising factors that might be associated with greater predominance of Enterobacterales and drug-resistant Enterobacterales, such as sink usage, ward-level antimicrobial usage, and patient populations, would also be of interest.

In conclusion, without conditioning on the presence of resistance markers, we have demonstrated that colonisation of ward sink drains with diverse and abundant populations of Enterobacterales, including drug-resistant strains, is common and persistent. The evidence linking contaminated, unmitigated wastewater reservoirs (including sink drains) in healthcare settings with outbreaks of colonisation/disease with drug-resistant Gram-negative bacilli in patients seems clear (5, 20), but no study to our knowledge has focused on the potential risk posed by Enterobacterales in sinks in general. Screening of sinks is not carried out in the absence of observed outbreaks, making it difficult to quantify wider patient-associated risk from the studies available. We demonstrate that contaminated sinks may be contributing to a proportion of healthcare-associated infections caused by Enterobacterales, and further work to investigate how to reduce the risk posed by this hospital environmental reservoir is warranted.

## MATERIALS AND METHODS

### Ward-based sink sampling

We sampled three units (acute admissions [AA], adult critical care [ACC], adult general medicine [female only] [GM]) within a single hospital (John Radcliffe Hospital, Oxford, UK) four times on rotation every three weeks over three months, March-May 2017. Units were chosen to capture different patient populations, admission turnaround times and wastewater plumbing infrastructure. The haematology ward (on a separate hospital site) was also sampled on a single day (12th May 2017) subsequent to a small cluster of patient cases of bla_OXA-48_ carbapenemase-associated Enterobacterales bloodstream infections [described previously (21). Ward and sink/wastewater layouts were obtained from estates, and each sink/drain site was assigned a unique site identifier (Table S1).

On each day of sampling, autoclaved tubing cut to 10 inches was used to aspirate from sink p-traps via a sterile 50ml syringe. Up to 50mls of fluid was aspirated where possible. 100μL of 10-fold dilutions (10^−2^, 10^−3^, 10^−4)^of each sink p-trap aspirate were plated onto CHROMagar Orientation media (Becton Dickinson, Franklin Lakes, NJ, USA), with no disc, cefpodoxime (10μg), ertapenem (10μg) (Thermo Scientific Oxoid, Basingstoke, UK) applied in a triangular fashion to each plate. Cultures were incubated at 37°C for ∼18hrs. Growth of Enterobacterales (presence/absence) and density (sparse/dense/confluent) in all zones was recorded (i.e. no antibiotic, in the presence of cefpodoxime, and in the presence of ertapenem). Up to four distinct colonies of each of presumptive *E. coli* and *Klebsiella* spp. were sub-cultured on CHROMagar Orientation to confirm purity and species identification. Species identification of sub-cultured colonies was confirmed by MALDI-ToF (MALDI Biotyper, Bruker, Billerica, MA, USA). Stocks of sub-cultured isolates were stored at −80°C in 400μl of nutrient broth + 10% glycerol prior to DNA extraction for sequencing. Aspirates from sink p-traps were then centrifuged at 4000 rpm for 10 minutes at 4°C, and supernatants removed; pellets were stored at −80°C.

### Patient isolate sampling

For AA, ACC, GM wards, a pseudo-anonymised, prospective feed was set-up to try and enable real-time capture of isolates from all samples culture-positive for *E. coli, K. pneumoniae* and *K. oxytoca* from patients that had been admitted to any of these wards during the study time period and were processed routinely through the clinical microbiology laboratory in the John Radcliffe Hospital in accordance with local standard operating procedures for clinical sample types, and compliant with national standards for microbiology investigations (22). These typically involve selective culture steps and species identification using MALDI-ToF (MALDI BioTyper, Bruker, Billerica, MA, USA).

Pseudo-anonymised extracts of all patient culture results and admission/discharge data covering the study period were obtained after the study was finished through the Infections in Oxfordshire Database (which has generic Research Ethics Committee, Health Research Authority and Confidentiality Advisory Group approvals [14/SC/1069, ECC5-017(A)/2009]) to enable an evaluation of *i)* baseline sampling denominators, *ii)* the extent of relevant clinical isolate capture, and *iii)* the temporal and spatial overlap of any genetically related sequenced isolates from patients and sequenced isolates/metagenomes from sinks.

### Isolate sequencing and p-trap aspirate metagenomics

All isolates confirmed as *E. coli, K. pneumoniae* and *K. oxytoca* from patients and p-trap aspirates were extracted for sequencing using the QuickGene DNA extraction kit (Autogen, MA, USA) as per the manufacturer’s instruction, plus an additional mechanical lysis step prior to chemical lysis (FastPrep, MP Biomedicals, CA, USA; 6m/s for two 40 second cycles).

For metagenomics, DNA was extracted from a subset of stored pellets (n=20) using the MoBio PowerSoil DNA isolation kit (Qiagen, Hilden, Germany) as per the manufacturer’s instructions, and including a mechanical lysis step of two 40 second cycles at 6m/s in lysing matrix E and final elution in buffer CDT-1 (Autogen, MA, USA). 45ng of Thermus thermophilus DNA (reference strain HB27, ATCC BAA-163 [DSMZ, Germany]) was added to each sample in the PowerBead tube at the start of the experiment, prior to the addition of solution C1 as an internal control and normalisation marker (12). Sink aspirates were selected for metagenomics sequencing to enable evaluation of *i)* microbiome differences within and between wards, *ii)* longitudinal change in microbiota composition, and *iii)* whether culture-negative sinks harboured the bacterial species being studies i.e. indicating limited sensitivity of culture-based approaches.

Short read sequencing (single isolate and metagenomics) was performed on the Illumina HiSeq 4000, pooling 192 isolate extracts and 6 metagenomes per lane, and generating 150bp paired-end reads.

### Computational methods

Cultured isolate informatics. Of the isolates sent for sequencing, 439/446 (98%) sink and 46/46 (100%) patient isolates were successfully sequenced and classified with Kraken/MiniKraken (23) as Enterobacterales, and used for subsequent analysis. Isolate consensus sequences were constructed by read mapping and consensus inference with respective *E. coli, K. oxytoca* and *K. pneumoniae* reference genomes AE014075.1, NC_018106.1 and CP000647.1 using Snippy 4.4.0 (24). Isolate genomes were assembled using Shovill 1.0.4 (25). Recombination-adjusted phylogenetic reconstruction was performed using runListCompare 0.3.8 (26) wrapping IQ-TREE 1.6.11 (27) and ClonalFrameML 1.12 (28). Final core genome alignments included 218/219 *E. coli* isolates, 165/167 *K. oxytoca* isolates and 98/99 *K. pneumoniae* isolates, all of which satisfied the runListCompare filtering criteria of perACGT_cutoff ≥70%, varsite_keep ≥0.8 and seq_keep≥0.7. 100 SNP core genome clusters were defined by single linkage clustering of runListCompare pairwise distance matrices. Trees were midpoint rooted prior to visualisation. See supplementary data repository for runListCompare configuration. Read-based MASH trees were constructed using MASH 2.2.2 (29) and RapidNJ 2.3.2 (30) using 21mers, a sketch size of 10,000 and a minimum abundance threshold of 10 *k*-mers. Assembly-based core and accessory genome partitioning was performed using PopPUNK 1.1.7 (31). Resistance genotyping and phenotype prediction in cultured isolates was performed using ResPipe and ARIBA 2.14.4 (32) with the CARD 3.0.3 database (33). Tree comparisons (tanglegrams) were generated using the R package Dendextend 1.5.0 (34).

### Metagenome informatics

Metagenomic sequences were analysed for taxonomic and antimicrobial resistance gene presence using ResPipe (12) and Kraken2 (35) with CARD database version 3.0.3. Large resistance gene families were clustered to facilitate visualisation of resistance profiles (Figure 4) (methodology documented in supplementary data repository). A metagenomic assembly of the *mcr-4* gene was generated with MEGAHIT 1.2.9 (36), to which reads were aligned with Minimap2 2.17-r941 (37) and consensus inferred using Kindel (38). Metagenomic summary statistics were generated using Pavian (39). Data analysis was performed with the SciPy ecosystem (40) and JupyterLab (41). Matplotlib (42), Bokeh and Microreact (43) were used for visualisation.

### Data availability

Raw sequencing data are available under NCBI SRA accessions PRJNA604910 and PRJNA604975 (cultured isolates), and ENA project PRJEB36775 (metagenomes). A supplementary data repository containing metadata, phylogenies, Jupyter notebooks, Microreact projects and Pavian reports is archived at https://figshare.com/articles/Enterobacterales_colonisation_of_hospital_sink_drains/11860893

## ACKNOWLEDGMENTS

This work was funded by the National Institute for Health Research Health Protection Research Unit (NIHR HPRU) in Healthcare Associated Infections and Antimicrobial Resistance at the University of Oxford, in partnership with Public Health England (PHE) [HPRU-2012-10041] and the NIHR Oxford Biomedical Research Centre. This work was supported by an Oxford University Clinical Academic Graduate School (OUCAGS) Career Development Support for Clinical Lecturers grant to NS. Extensive use was made of The Oxford Biomedical Research Computing (BMRC) facility, a joint development between the Wellcome Centre for Human Genetics and the Big Data Institute supported by Health Data Research UK and the NIHR Oxford Biomedical Research Centre. Financial support was also provided by the Wellcome Trust Core Award Grant Number 203141/Z/16/Z. NS is funded by a University of Oxford/Public Health England lectureship. TEAP and ASW are NIHR Senior Investigators. The views expressed are those of the author(s) and not necessarily those of the NHS, the NIHR or the Department of Health.

We thank the staff of the microbiology laboratory at OUH NHS Foundation Trust, the healthcare staff on the participating units, and Neil Hemmings and the Estates team at OUH for their cooperation in sample collection. Valuable discussion with Timothy Davies, Liam Shaw and Jeremy Swann informed computational analysis.

## SUPPLEMENTARY MATERIAL

**Table S1**. Surveyed sink descriptions and *E. coli*/*Klebsiella* spp. culture results across timepoints.

**Table S2**. Cultured Enterobacterales species by ward.

**Table S3**. Pairwise Mantel correlation of different within-species distance matrices. These include recombination-adjusted core SNP phylogeny (reads-core-snp), read-based MASH distance (reads-mash) and PopPUNK estimates of core and accessory genomic distance from de novo assemblies (assemblies-core-mash, assemblies-accessory-mash).

**Table S4**. Permutational analysis of variance. Permutation tests for association of genetic structure with ward (n=3) and sink (n=18) for three species of sink drain Enterobacterales. Corresponding test results are shown for differential dispersion between groups (PERMDISP). Bold type indicates significant (p<0.05) group association under PERMANOVA in the absence of significant differential dispersion (PERMDISP).

**Table S5**. *mcr-4* coverage. Sequencing coverage and mean depth of the 1,626bp metagenome-assembled *mcr-4* gene from sink A10, to which metagenomic short reads mapped from three sinks (including A10) across six sink-timepoints within the general medicine ward.

**Figure S1**. Cultured strains observed on the Haematology ward. Different colours indicate distinct 100 core SNP strains, and cefpodoxime-resistant and/or ESBL gene-positive isolates are indicated by filled markers.

**Figure S2**. Spatial structure of sink metagenome *k*-mer composition. Left and centre: visualisation of 31mer pairwise MASH distances of total metagenome content using hierarchical clustering (left) and multidimensional scaling (centre). Right: comparison of within sink, within ward and between ward pairwise MASH distances.

**Figure S3**. Receiver operating characteristic (ROC) for detection of Enterobacterales by culture with varying metagenomic abundance.

**Figure S4**. Tanglegrams comparing recombination-corrected core phylogenies and read-based whole genome MASH + neighbour joining phylogenies for a) *E. coli*, b) *K. oxytoca* and c) *K. pneumoniae*. Topologically consistent subtrees are rendered with solid branches.

**Figure S5**. *Klebsiella* spp. lineage persistence in cultured sink drain aspirates and contemporaneous clinical isolates from patients with ward contact during the sampling period.

