## Supplementary tables and figures for "Multi-omic surveillance of *Escherichia coli* and *Klebsiella* spp. in hospital sink drains and patients"

| | | | Timepoint | | | |
|---|---|---|---|---|---|---|
| Sink | Ward | Sink location | 1 | 2 | 3 | 4 |
| A1 | General medicine | Staff toilet (entrance) | positive | positive | positive | positive |
| A3 | General medicine | Patient sideroom 1 (1 bed) | negative | positive | positive | negative |
| A4 | General medicine | Patient sideroom 2 (1 bed) | negative | positive | positive | positive |
| A5 | General medicine | Patient toilet (entrance) | negative | negative | negative | negative |
| A7 | General medicine | Patient bay 1 (4 beds) | positive | positive | positive | negative |
| A8 | General medicine | Patient bay 2 (4 beds) | negative | positive | positive | positive |
| A9 | General medicine | Patient bay 3 (4 beds) | positive | positive | positive | positive |
| A10 | General medicine | Patient bay 4 (4 beds) | positive | positive | positive | positive |
| A11 | General medicine | Patient toilet | negative | negative | negative | negative |
| A13 | General medicine | Patient sideroom 3 (1 bed) | negative | negative | negative | positive |
| A14 | General medicine | Patient sideroom 4 (1 bed) | positive | positive | positive | positive |
| A15 | General medicine | Patient toilet 1 | negative | negative | negative | negative |
| A18 | General medicine | Patient toilet 2 | negative | negative | negative | negative |
| A21 | General medicine | Patient toilet 3 | negative | negative | negative | negative |
| A23 | General medicine | Medicine preparation room | negative | negative | negative | negative |
| A24 | General medicine | Reception area | negative | positive | positive | negative |
| A25 | General medicine | Staff room | positive | positive | positive | positive |
| A26 | General medicine | Sluice room | negative | negative | negative | negative |
| A27 | General medicine | Sluice room | negative | negative | negative | negative |
| A28 | General medicine | Patient room sideroom 5 (1 bed) | negative | positive | positive | positive |
| B1 | Acute critical care | Relatives' day room | positive | positive | positive | positive |
| B2 | Acute critical care | Relatives' toilet | negative | negative | negative | negative |
| B3 | Acute critical care | Staff toilet M | negative | negative | positive | negative |
| B4 | Acute critical care | Staff toilet F | negative | negative | negative | negative |
| B5 | Acute critical care | Patient sideroom 1 (1 bed) | negative | negative | positive | negative |
| B6 | Acute critical care | Patient bay 1 (6 beds) | negative | negative | negative | negative |
| B7 | Acute critical care | Patient sideroom 2 (1 bed) | negative | negative | negative | positive |
| B8 | Acute critical care | Patient bay 1 (6 beds) | negative | negative | negative | negative |
| B9 | Acute critical care | Patient bay 1 (6 beds) | positive | negative | negative | positive |
| B10 | Acute critical care | Laboratory | negative | negative | negative | negative |
| B11 | Acute critical care | Laboratory | negative | negative | negative | negative |
| B12 | Acute critical care | Staff room kitchen sink | positive | positive | positive | positive |
| B13 | Acute critical care | Sluice room | negative | negative | negative | negative |
| B14 | Acute critical care | Sluice room | negative | negative | negative | negative |
| B15 | Acute critical care | Sluice room | negative | negative | negative | negative |
| B16 | Acute critical care | Patient bay 2 (4 beds) | negative | negative | positive | negative |
| B17 | Acute critical care | Patient bay 2 (4 beds) | negative | negative | positive | positive |
| B18 | Acute critical care | Patient bay 2 (4 beds) | negative | negative | negative | negative |
| B19 | Acute critical care | Patient bay 3 (4 beds) | negative | positive | positive | positive |
| B20 | Acute critical care | Patient bay 3 (4 beds) | negative | negative | negative | negative |
| B21 | Acute critical care | Patient sideroom 3 (1 bed) | positive | negative | positive | negative |
| B22 | Acute critical care | Patient sideroom 4 (1 bed) | negative | negative | negative | negative |
| B23 | Acute critical care | Dirty utility sink | negative | negative | negative | negative |
| C1 | Acute admissions | Staff toilet | negative | positive | positive | negative |
| C2 | Acute admissions | Patient sideroom 1 (1 bed) | positive | positive | positive | positive |
| C3 | Acute admissions | Female patient toilet | negative | positive | negative | negative |
| C4 | Acute admissions | Female patient toilet and shower | negative | negative | negative | negative |
| C5 | Acute admissions | Patient bay 1 (6 beds) | positive | positive | negative | positive |
| C6 | Acute admissions | Patient bay 2 (4 beds) | positive | positive | positive | positive |
| C7 | Acute admissions | Patient sideroom 2 (1 bed) | negative | positive | negative | negative |
| C8 | Acute admissions | Patient sideroom 3 (1 bed) | negative | positive | negative | positive |
| C9 | Acute admissions | Patient sideroom 4 (1 bed) | positive | negative | positive | positive |
| C10 | Acute admissions | Patient bay 3 (4 beds) | positive | positive | positive | positive |
| C11 | Acute admissions | Patient bay 4 (4 beds) | positive | negative | negative | positive |
| C12 | Acute admissions | Patient sideroom 5 (1 bed) | negative | positive | negative | negative |
| C13 | Acute admissions | Patient sideroom 6 (1 bed) | negative | negative | negative | negative |
| C14 | Acute admissions | Patient bay 5 (7 beds) | positive | positive | positive | positive |
| C15 | Acute admissions | Patient bay 5 (7 beds) | negative | positive | positive | negative |
| C16 | Acute admissions | Near-patient testing room | positive | positive | positive | negative |
| C17 | Acute admissions | Patient toilet waiting room | negative | positive | negative | negative |
| C18 | Acute admissions | Ambulatory bay | positive | positive | negative | negative |
| C19 | Acute admissions | Male patient toilet | negative | negative | negative | negative |
| C20 | Acute admissions | Male patient toilet and shower | negative | negative | negative | negative |
| C21 | Acute admissions | Dirty utility sink | negative | negative | negative | negative |
| C22 | Acute admissions | Patient toilet | negative | negative | negative | negative |
| C23 | Acute admissions | Treatment/medicines room | negative | negative | negative | negative |

**TABLE S1.** Surveyed sinks

| ward | species | count | ward sum | % |
|---|---|---|---|---|
| GM | *E. coli* | 59 | 179 | 33% |
| | *K. oxytoca* | 56 | 179 | 31% |
| | *K. pneumoniae* | 64 | 179 | 36% |
| ACC | *E. coli* | 36 | 64 | 56% |
| | *K. oxytoca* | 10 | 64 | 16% |
| | *K. pneumoniae* | 18 | 64 | 28% |
| AA | *E. coli* | 79 | 166 | 48% |
| | *K. oxytoca* | 76 | 166 | 46% |
| | *K. pneumoniae* | 11 | 166 | 7% |
| HAEM | *E. coli* | 6 | 30 | 20% |
| | *K. oxytoca* | 24 | 30 | 80% |
| | *K. pneumoniae* | 0 | 30 | 0% |

**TABLE S2.** Cultured Enterobacterales by ward. The distribution of cultured Enterobacterales target species by ward

| Species | Pairwise distance comparison | | Mantel *r* | *p* |
|---|---|---|---|---|
| *E. coli* | reads-core-snp | reads-mash | 0.879 | 0.001 |
| | | assemblies-core-mash | 0.987 | 0.001 |
| | | assemblies-accessory-mash | 0.892 | 0.001 |
| | reads-mash | assemblies-core-mash | 0.893 | 0.001 |
| | | assemblies-accessory-mash | 0.870 | 0.001 |
| | assemblies-core-mash | assemblies-accessory-mash | 0.933 | 0.001 |
| *K. oxytoca* | reads-core-snp | reads-mash | 0.865 | 0.001 |
| | | assemblies-core-mash | 0.901 | 0.001 |
| | | assemblies-accessory-mash | 0.857 | 0.001 |
| | reads-mash | assemblies-core-mash | 0.995 | 0.001 |
| | | assemblies-accessory-mash | 0.951 | 0.001 |
| | assemblies-core-mash | assemblies-accessory-mash | 0.938 | 0.001 |
| *K. pneumoniae* | reads-core-snp | reads-mash | 0.808 | 0.001 |
| | | assemblies-core-mash | 0.800 | 0.001 |
| | | assemblies-accessory-mash | 0.891 | 0.001 |
| | reads-mash | assemblies-core-mash | 0.996 | 0.001 |
| | | assemblies-accessory-mash | 0.942 | 0.001 |
| | assemblies-core-mash | assemblies-accessory-mash | 0.916 | 0.001 |

**TABLE S3.** Pairwise Mantel correlation of different within-species distance matrices. These include recombination-adjusted core SNP phylogeny (reads-core-snp), read-based MASH distance (reads-mash) and PopPUNK estimates of core and accessory genomic distance from *de novo* assemblies (assemblies-core-mash, assemblies-accessory-mash).

| | | | PERMANOVA | | PERMDISP | |
|---|---|---|---|---|---|---|
| Factor | Species | Distances | Pseudo-F | *p* | Pseudo-F | *p* |
| Sink | *E. coli* | assemblies-acc-mash | 11.74 | 0.001 | 11.67 | 0.001 |
| | | assemblies-core-mash | 8.31 | 0.001 | 10.52 | 0.001 |
| | | core-snp | 7.21 | 0.001 | 5.68 | 0.001 |
| | | reads-mash | 6.30 | 0.001 | 5.45 | 0.001 |
| | *K. oxytoca* | assemblies-acc-mash | 9.61 | 0.001 | 3.05 | 0.001 |
| | | assemblies-core-mash | 10.62 | 0.001 | 2.53 | 0.001 |
| | | core-snp | 12.85 | 0.001 | 2.01 | 0.001 |
| | | reads-mash | 9.53 | 0.001 | 2.66 | 0.001 |
| | ***K. pneumoniae*** | **assemblies-acc-mash** | **13.03** | 0.001 | 1.56 | 0.133 |
| | | **assemblies-core-mash** | **9.65** | 0.001 | 1.49 | 0.084 |
| | | **core-snp** | **9.51** | 0.001 | 1.38 | 0.337 |
| | | **reads-mash** | **10.30** | 0.001 | 1.54 | 0.078 |
| Ward | *E. coli* | assemblies-acc-mash | 20.98 | 0.001 | 23.16 | 0.001 |
| | | assemblies-core-mash | 18.16 | 0.001 | 29.86 | 0.001 |
| | | core-snp | 25.48 | 0.001 | 51.71 | 0.001 |
| | | reads-mash | 12.57 | 0.001 | 12.45 | 0.001 |
| | *K. oxytoca* | assemblies-acc-mash | 18.71 | 0.001 | 47.19 | 0.001 |
| | | assemblies-core-mash | 21.54 | 0.001 | 86.31 | 0.001 |
| | | core-snp | 24.59 | 0.001 | 43.62 | 0.001 |
| | | reads-mash | 20.86 | 0.001 | 83.74 | 0.001 |
| | ***K. pneumoniae*** | **assemblies-acc-mash** | **12.15** | 0.001 | 0.45 | 0.656 |
| | | **assemblies-core-mash** | **12.40** | 0.001 | 1.17 | 0.307 |
| | | **core-snp** | **7.53** | 0.001 | 0.99 | 0.407 |
| | | **reads-mash** | **13.32** | 0.001 | 1.62 | 0.185 |

**TABLE S4.** Permutational analysis of variance. Permutation tests for association of genetic structure with ward (n=3) and sink (n=18) for three species of sink drain Enterobacterales. Corresponding test results are shown for differential dispersion between groups (PERMDISP). Bold type indicates significant ($p<0.05$) group association under PERMANOVA in the absence of significant differential dispersion (PERMDISP).

| | *mcr-4* gene coverage (%) | mean depth |
|---|---|---|
| sink-timepoint | | |
| A10T1 | 100.0 | 31.6 |
| A10T4 | 92.4 | 2.7 |
| A8T1 | 73.1 | 1.1 |
| A8T4 | 14.2 | 0.2 |
| A9T1 | 51.5 | 0.6 |
| A9T4 | 32.2 | 0.4 |

**TABLE S5.** *mcr-4* coverage. Sequencing coverage and mean depth of the 1,626bp metagenome-assembled *mcr-4* gene from sink A10, to which metagenomic short reads mapped from three sinks (including A10) across six sink-timepoints within the general medicine ward.

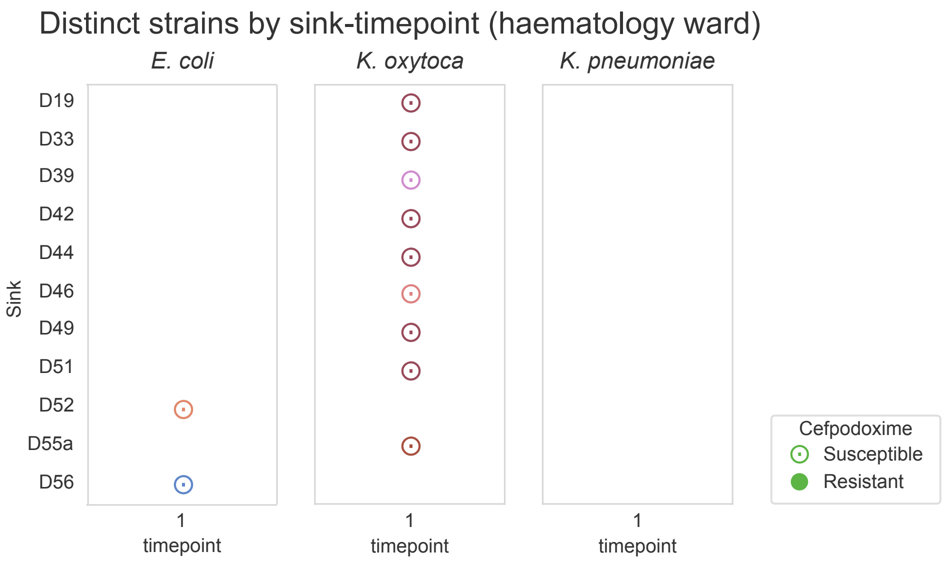


**FIGURE S1. Cultured strains observed on the haematology ward. Different colours indicate distinct 100 core SNP strains, and cefpodoxime-resistant and/or ESBL gene-positive isolates are indicated by filled markers.**

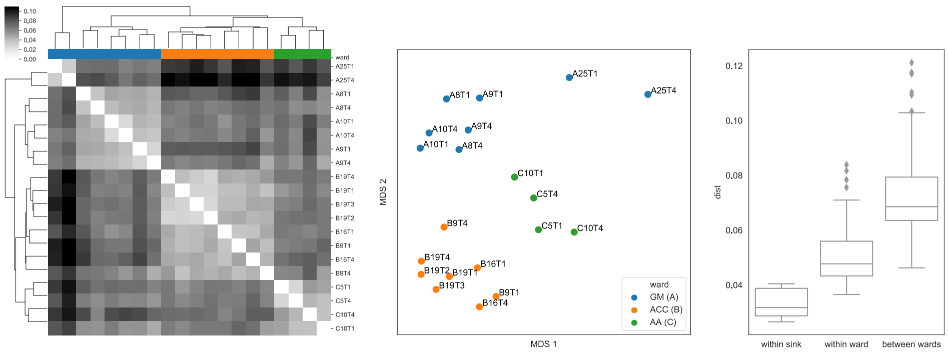


**FIGURE S2. Spatial structure of sink metagenome *k*-mer composition. Left and centre: visualisation of 31mer pairwise MASH distances of total metagenome content using hierarchical clustering (left) and multidimensional scaling (centre). Right: comparison of within sink, within ward and between ward pairwise MASH distances.**

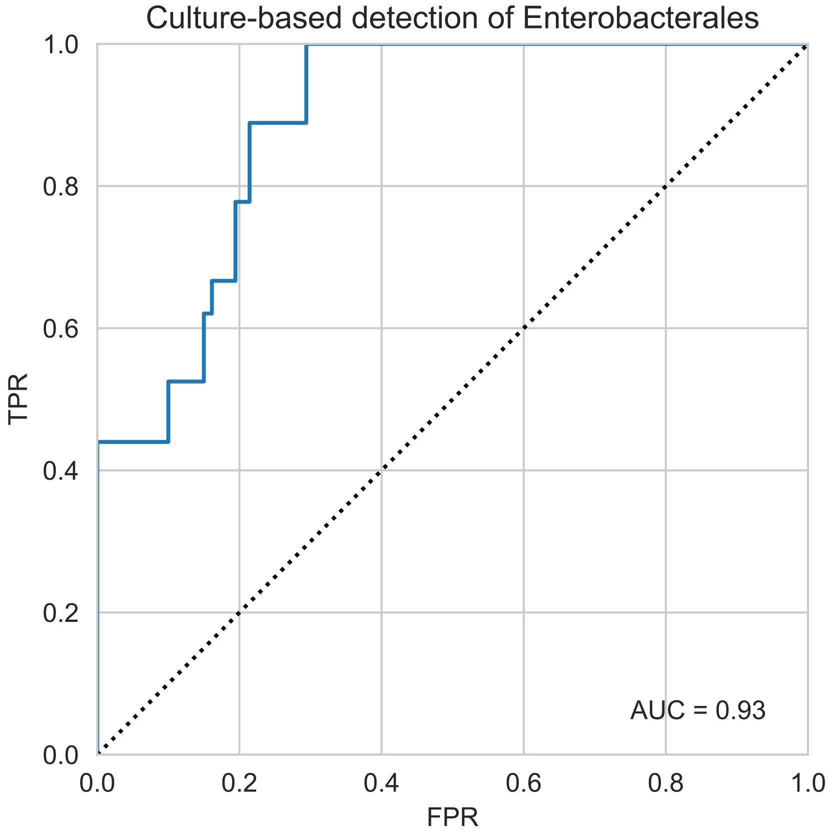


**FIGURE S3. Receiver operating characteristic (ROC) for detection of Enterobacterales by culture with varying metagenomic abundance.**

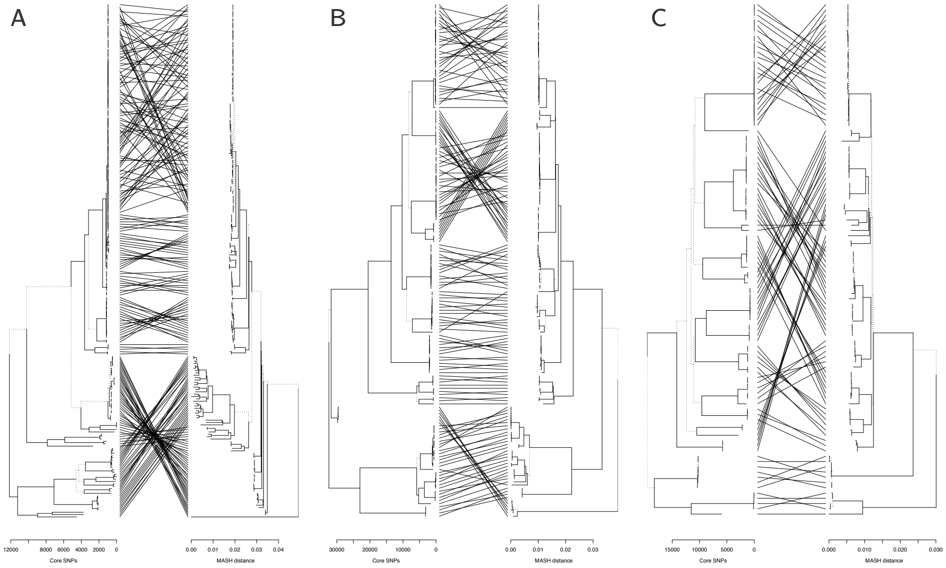


**FIGURE S4. Tanglegrams comparing recombination-corrected core phylogenies and read-based whole genome MASH + neighbour joining phylogenies for *a) E. coli*, *b) K. oxytoca* and *c) K. pneumoniae*. Topologically consistent subtrees are rendered with solid branches.**

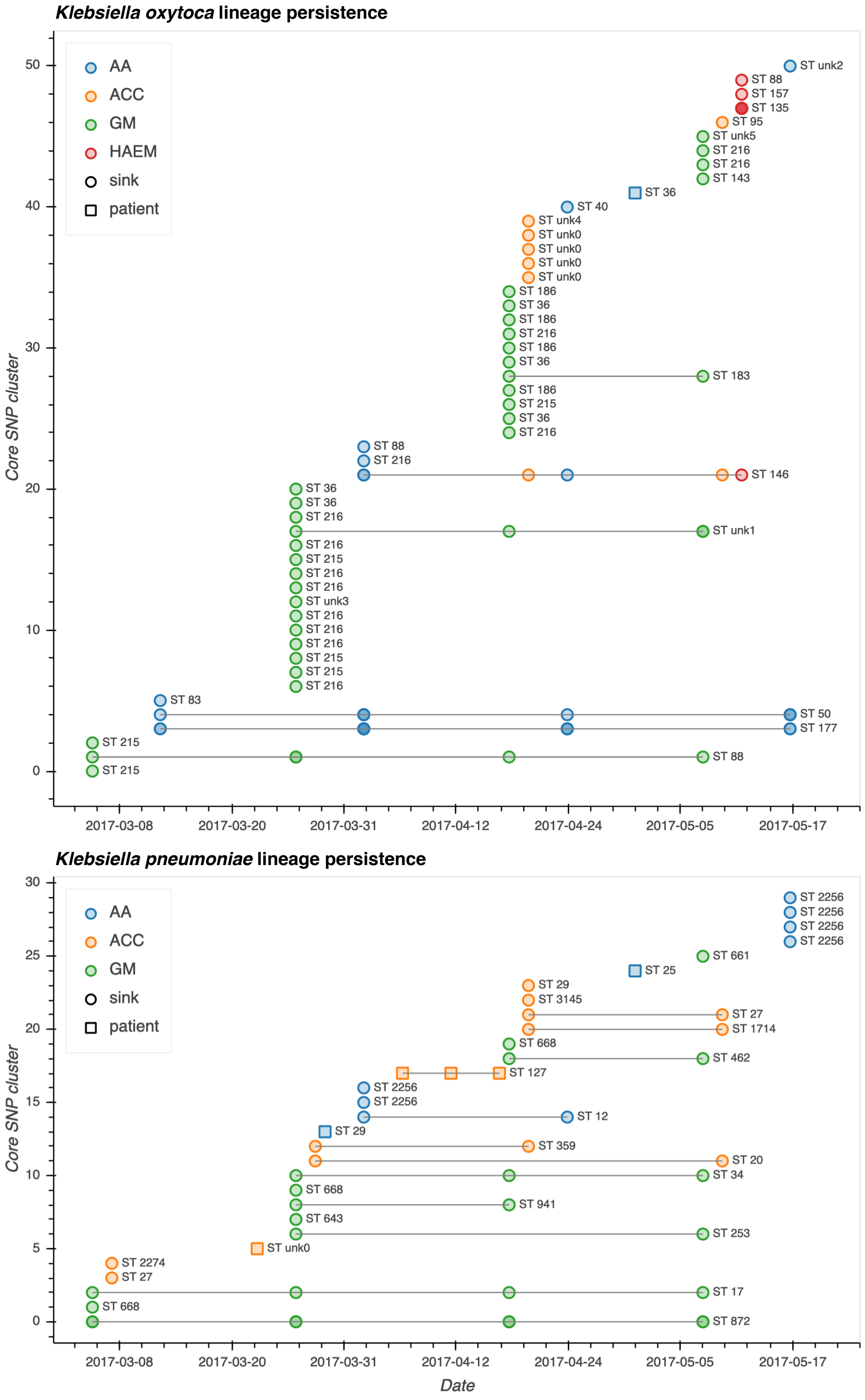


**FIGURE S5.** ***Klebsiella*** **spp. lineage persistence in cultured sink drain aspirates and contemporaneous clinical isolates from patients with ward contact during the sampling period.**
